# The memory of hyperosmotic stress response in yeast is modulated by gene-positioning

**DOI:** 10.1101/625756

**Authors:** Zacchari Ben Meriem, Yasmine Khalil, Pascal Hersen, Emmanuelle Fabre

## Abstract

Cellular memory is a critical ability displayed by microorganisms in order to adapt to potentially detrimental environmental fluctuations. In the unicellular eukaryote *S. cerevisiae* cellular memory can take the form of a faster or a decreased response following repeated stresses in cell population. Using microfluidics and fluorescence time-lapse microscopy, we studied how yeasts respond to short-pulsed hyperosmotic stresses at the single-cell level by analyzing the dynamical behavior of the stress responsive STL1 promoter fused to a fluorescent reporter. We established that pSTL1 shows variability in its successive activations following two repeated short stresses. Despite this variability, most cells displayed a memory of past stresses through a decreased activity of pSTL1 upon repeated stress. Notably, we showed that genomic location is important for the memory effect since promoter displacement to a pericentromeric chromatin domain leads to a decreased transcriptional strength of pSTL1 and to the loss of memory. This study provides a quantitative description of a cellular memory that includes single-cell variability and points towards the contribution of the chromatin structure in stress memory.

## Introduction

Cellular memory can be defined as a cellular response to transient and repeated stimuli. These latter can emanate from constantly fluctuating and potentially stressful environments, thus possibly exerting a selective pressure on cell viability [1]. To ensure their survival, living organisms have developed various strategies to cope with environmental changes. One possible way for cells to maintain their biological functions in a challenged environment is to regulate gene transcription [2]. The active genetic response allowing cells to survive a single stimulus is referred to as cellular adaptation. Factors such as histone post-translational modifications, chromatin remodelers, specific proteins produced during the stress or even changes in chromatin conformation have been determined to be causal factors for adaptation to environmental changes [3, 4]. What happens when cells encounter consecutive stresses is less well understood, however it has been observed in some cases that the adaptation to a first stress can serve as a learning process to a better adaptation to a consecutive stress. This process is defined as memory. Biological mechanisms known to underlie cellular memory involve chromatin remodeling (epigenetic memory) or proteins synthesized during the first stress, which might leave a trace of the first event (cytoplasmic memory) [5–7].

In the budding yeast, studies on cellular memory showed that cells confronted to successive environmental stresses can respond differently. For instance, the so called galactose memory is characterized by a faster transcriptional reactivation of the GAL cluster, while an example of the memory of long hyperosmotic stresses is characterized by a reduced activity of the osmo-responsive gene *GRE2* without any difference in the time of reactivation of this gene [5, 7].

All eukaryotes have a highly organized nucleus. Yeast chromosomes follow a Rabl organization; centromeres are tethered to the spindle pole body, while telomeres are anchored to the nuclear periphery [8, 9]. Interestingly, the galactose or inositol memory appears to rely on 3D gene positioning, since repositioning of the *INO1* gene or *GAL* cluster towards the nuclear periphery, in an H2AZ and nucleoporin-dependent manner, is important for memory [7, 10, 11]. The nuclear organization may also play a critical role in stress response as most stress response genes are located in subtelomeres. Subtelomeres lack essential genes but are enriched in fast-evolving non-essential gene families that are needed to adapt to environmental changes [12]. Subtelomeres are subjected to silencing by proteins of the Silent Information Regulator (SIR) complex, but stress conditions can lift this repression [13, 14].

Most studies questioning memory effects are carried out on isogenic populations of cells, giving information on the mean behavior of the population [15]. Nevertheless, cells populations are heterogeneous due to extrinsic noise, such as age, size or position in the cell cycle (for reviews, [16, 17]). Moreover gene expression is an inherently stochastic phenomenon, because of the low number and availability of transcription factors, accessibility of the promoter or functional regulatory networks [18]. Overall, stochasticity causes genetically identical cells to exhibit different behavior when encountering the same stimuli.

The response to osmotic changes in the budding yeast has proven to be a good tool to study the emergence of adaptation and cellular memories in this organism [19, 20]. When yeast faces an osmolarity increase in its environment (hyperosmotic stress), intracellular water flows out of the cell, leading to its shrinkage [21]. The imbalance of osmotic pressure is detected by osmosensors that activate the High Osmolarity Glycerol (HOG) pathway that phosphorylates the cytoplasmic protein Hog1 [22, 23]. Phosphorylated Hog1 translocates into the nucleus where it participates in the activation and regulation of an estimated 10% of the genome, including the osmo-responsive gene STL1 [24]. Thanks to the HOG pathway, yeast can physiologically adapt to a hyperosmotic stress in 15-30 min [25], notably by producing glycerol, which leads to homeostasis. The dephosphorylation of Hog1 and its exit from the nucleus signals the end of the adaptation to the hyperosmotic stress.

We here present a single-cell study of *S. cerevisi*ae cells submitted to short pulsed hyperosmotic stresses in a well-controlled system based on time lapse fluorescence microscopy and microfluidics [26, 27]. Hundreds of single cells receiving repeated osmotic stresses were tracked and analyzed. We found that in response to two consecutive hyperosmotic stresses separated by 4h, individual cells display variability in the dynamical activity of pSTL1 in response to the two stresses. Despite the existence of this pronounced dynamical variability, most of the cells adopt the same behavior, consisting in decreased amplitude of response upon stress. We called this specific behavior memory effect. Importantly, we found that the chromatin environment modulates cells response to pulsed stresses. Relocation of the promoter of interest close to the centromere causes a reduced activity of pSTL1 and loss of the memory effect. Overall, our study suggests that the specific location of pSTL1 at the subtelomere is required for an optimal level of transcription that can go beyond a simple stochastic behavior and lead to the emergence of a memory in response to short osmotic stresses.

## Results

### The response of a population to successive hyperosmotic stresses suggests cellular memory

A population of growing yeast cells in a microfluidic device was submitted to short and repeated hyperosmotic stresses, using 1M Sorbitol (figure 1a). To measure the response to hyperosmotic stress, we used a reporter of the activity of the HOG pathway in which pSTL1 promoter is tagged with the yECITRINE fluorescent protein (yEFP) allowing for fluorescence quantification at the single cell level as a function of time [28] (figure 1b). During the time course of an experiment, each cell is tracked allowing for quantification of the first and second activation of pSTL1 due to the short and transient activation of the HOG pathway (figure 1c). The limited duration of the stress (8 min) and the long delay between the two stresses (4 hours) guaranteed i) investigation on the genetic response to hyperosmotic stress before adaptation is established, 15-30 min after stress [29] and ii) full recovery of the cells from the first stress, thus allowing to compare the dynamics of response to the first and second osmotic stress. Moreover daughter cells born between the two stresses were not considered, since they did not receive the first stress and may blur the stress response. Rather, we focus exclusively on the population that received both the first and the second stress (figure 1c). In such a population, we calculated the mean pick of fluorescence reached during the first and second stress, taking the basal fluorescence level prior to the corresponding stress as a reference. It reaches 77,14 ± 6,71 (a.u.) after the first stress and 55,44 ± 4,26 (a.u.)after the second stress, indicating a decrease of fluorescence intensity by 20% in average after the second stress (figure 1d). No difference in the time required to reach the peak of fluorescence was detected between the two picks (figure 1e). The decrease of fluorescence amplitude correlated with decreased protein amount, as detected by Western Blotting (supplementary figure 1), suggesting it is independent of photo bleaching.

**Figure 1.**
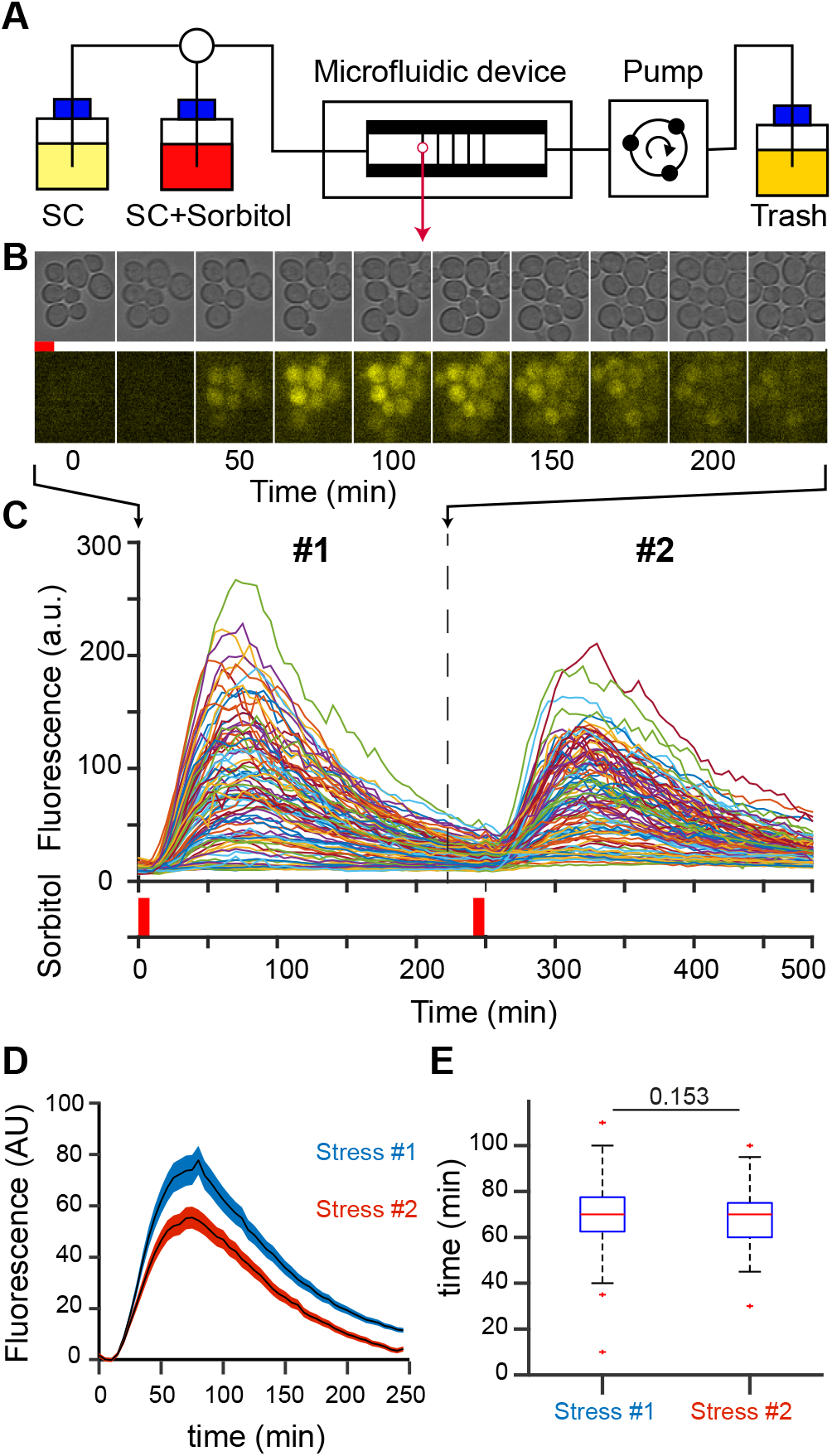
(A) Experimental setup. We use a multi-layer H-shaped microfluidics device composed of two large flow channels of 50µm height and 40µm thin, observation chambers of 400µm × 400µm × 3.7µm. Cells are trapped in the chambers and grow as a monolayer, facilitating cell segmentation and tracking. The medium flowing in the channels diffuse in the chambers. The hyperosmotic stresses activating expression of pSTL1-yECRITRINE are triggered using sorbitol 1M for 8min. SC, synthetic complete medium. (B) Example of cells in the microfluid chamber, submitted to an 8min stress and left to recover for 4h (240min.). Cells are imaged every 5min in bright light (20ms exposure, upper raw) and fluorescence light (200ms exposure, lower raw). (C) Fluorescent signal of Individual cells exposed to two successive 8 minutes stress separated by 4 hours. Hyperosmotic stress is schematized by red bars. (D) Fluorescence response of a population of N= 97 cells. The mean response of cells submitted to a first stress (dark blue), followed by another stress 4h later (light blue) is represented with the standard error of the mean. Fluorescence levels are normalized by the value of the peak of fluorescence during the first stress. The peak of fluorescence decreases upon stress. (E) Analyses of time response to two consecutive stresses in the same population. Time between stress induction and fluorescence peak reveals that the time of response is similar upon stress.

Altogether, these observations suggest that at the level of stressed cells population, there is a memory of the first stress event. Moreover, the decrease of fluorescence intensity at the second stress is seemingly due to a reduction of protein production rate rather than a shortened duration of transcription events.

### At the level of single cells, most cells, but not all, show a cellular memory

Yet, this memory effect was not shared equally among cells. Indeed, single-cell analysis reveals a dynamic variability in the individual response to the repeated stresses. We classified single-cell fluorescent trajectories according to typical behaviors based on the first and second responses to stress (figure 2a). The most frequent behavior (55% ± 11%) is in line with the population memory effect in which the cells submitted to the second stress showed lower fluorescence intensity (figure 2b). However, up to 18%±7% of cells display an opposite behavior, with a stronger response at the second stress (figure 2b). Very few cells showed similar responses to both stresses. Interestingly, we observed two subpopulations of cells that did not respond to one of the stress (figure 2b), although we confirmed that these cells indeed perceived the stress by visualization of transient cell shrinkage upon stress (Supplementary video & supplementary figure 2). Altogether our results show that the population behavior hides a richer set of dynamic behaviors of single-cell responses, which are likely the traces of the variability of the activation of pSTL1 by a hyperosmotic stress.

**Figure 2.**
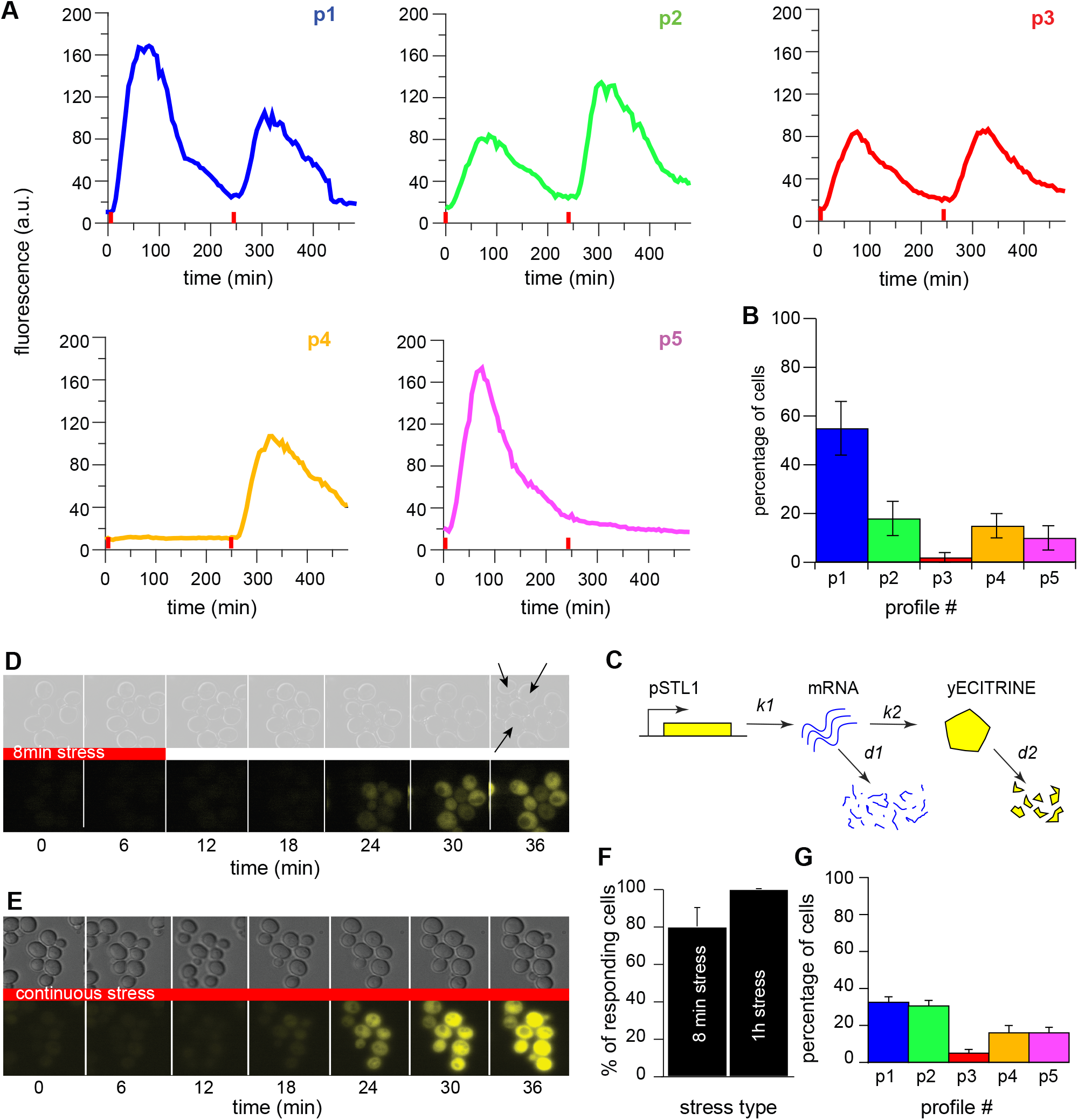
(A) Examples of five typical single-cell response profiles. Although the single-cell analysis reveals a dynamical variability of the response, we have defined five typical profiles of response adopted by the cells (p1 to p5). (B) Single-cell clustering. Using criteria on the values of the fluorescence peaks, dynamical variability of response was clustered according to the five typical fluorescence response. Errors bars represent standard errors. 708 cells were analyzed from 3 independent experiments. (C) Modeling gene expression upon stress in a memory-free system. Stochastic simulations with Gillespie algorithm were used to model the transcription of the fluorescent reporter and the protein translation upon stress. (D) Sequence images of cells submitted to an 8min stress. The arrows on the last bright field image show the cells that do not respond to the stress. (E) Sequence images of cells submitted to a continuous stress. All cells show a response to such a stress. (F) Quantification of responsive cells. Upon receiving an 8min stress, 80% of cells show a response, whereas 100% of cells respond to a 1h stress. (G) Single-cell quantification of computed cells according to the five typical profiles. The model in this case includes a transcription delay randomly chosen between 0 and 10min for each computed cells. This delay is also different during the two stresses. The simulation is run twice and the fluorescence peaks values are used to cluster the computed cells responses according to the five typical profiles of response.

### Cellular memory overcomes the stochasticity of gene expression on average

To determine the importance of the intrinsic variability on setting the different dynamical behaviors shown in figure 2a, that is the stochastic nature of the pSTL1 activation, we performed stochastic simulations based on the Gillespie algorithm [30]. We simulated the transcription of pSTL1 and the translation of the fluorescent reporter of 1000 cells submitted to two 8min stresses separated by 4h (figure 2c). The rates of production and degradation of mRNA and proteins were set as established previously [31] (supplementary Table 1). Such a model implies that the computed cells will necessarily respond to both stresses. However, our experiments show that cells exposed to an 8min stress do not necessarily respond to a stress (figure 2d), conversely to cells submitted to a longer stress (figure 2e). More specifically, the absence of response disappears for longer stress duration since for a 1h stress 100% cells showed an activation of pSTL1, while for 8 min stress 80% of cells showed a response (figure 2f). This experimental observation suggests the existence of a critical time from which all cells will eventually respond to a stress. A stochastic time of activation of the STL1 promoter was therefore added to the model. As expected in such a memory-free system, cells were clustered equally in the two main categories obtained experimentally, and the population did not display any memory effect. Of note, transcriptional delay made possible clusters 4 and 5 appearance (figure 2g).

The differences in clusters observed in vivo and through simulation suggest that a biological mechanism other than noise in transcription and translation is at play in the memory effect.

### Memory to pulsed hyperosmotic stress does not require de novo protein synthesis during the stress

In order to investigate the biological origin of the memory effect, we first wanted to determine if the memory effect was linked to one or several long-lived proteins synthesized during the first episode of stress. To test this hypothesis, we inhibited transcription upon stress using thiolutin, a well-studied molecule that inhibits all three RNA polymerases in the yeast in a reversible manner [32] (figure 3a, b). As expected, treatment of cells with thiolutin led to the loss of pSTL1 activity: when cells were treated for 1h with thiolutin (50µg/µL), no cells showed a fluorescent signal when stimulated by an 8 min hyperosmotic stress, while 80% ± 20% cells showed a signal in response to a hyperosmotic stress in the absence of thiolutin (figure 3a, b). We next tested the cells ability to respond back to a hyper-osmotic stress after treatment with thiolutin (figure 3c, d). After 1h in the presence of thiolutin, the inhibitor was washed out and cells were submitted to an 8 min hyperosmotic stress, 4h later. At the population level, a response similar to the response to the first stress without thiolutin treatment was observed (figure 3c, d). In the presence of thioluthin, stress response is however slightly decreased and the maximum of intensity after 8 min of stress is 88,55 +/− 7,7%, as compared to 100 +/− 4,74 % in non-treated cells, suggesting that not all of the thioluthin effect is erased. We then performed a thiolutin treatment during 1h, including during the first stress, then washed the inhibitor and submitted cells to a second stress 4h later (figure 3e, f). In these conditions, cells showed a marked decrease of the YFP signal by 40%, comparable to the 47% decreased response to a second stress without thiolutin treatment. Consequently, this experiment suggests that the memory effect is not primarily driven by de novo synthesis of proteins during the first stress, which could help the cells to respond to the second stress. To explain the observed memory effect, we can hypothesize that the first stress induces chromatin modifications independently of the activity of any RNA polymerases, but with an effect on subsequent transcription events at the pSTL1 locus. A possibility could be that chromatin marks would appear in most cells during the first stress and alter the response dynamics of cells during the second stress.

**Figure 3.**
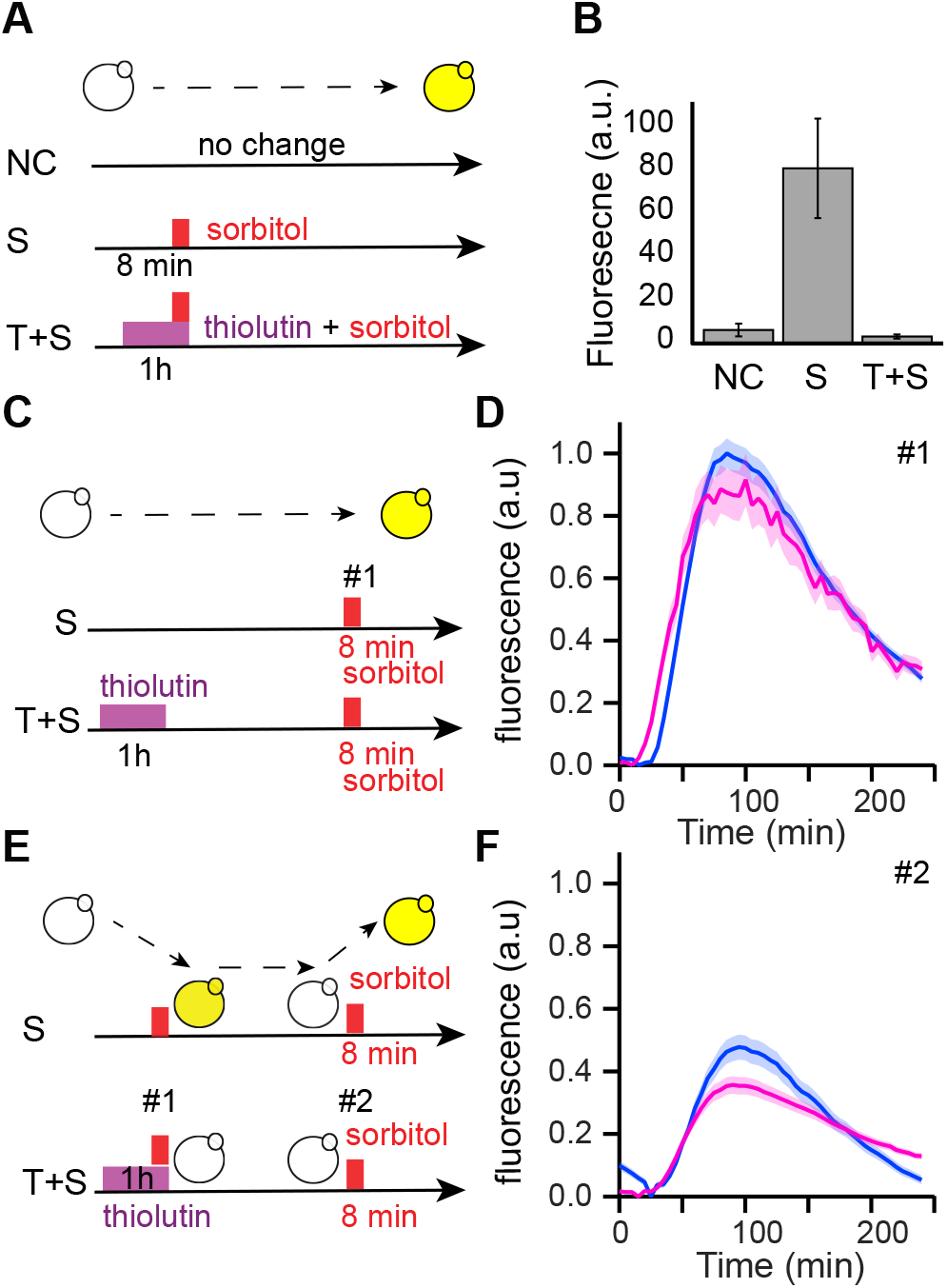
(A) Principle of the transcription inhibition experiment with thioluthin. A 1h thiolutin treatment (pink rectangle)) is performed prior and during the stress (red rectangle) in order to prevent the activity of pSTL1 upon stress. Non-stressed cells received no sorbitol (NC) and stressed cells received an 8 min hyperosmotic stress (red rectangle, S). (B) Single-cell quantification of the fluorescence of cells after 70 min in non-stress conditions (NC), hyperosmotic conditions (S) and hyperosmotic conditions in the presence of thiolutin (T + SC). (C) Principle of the experiment controlling thiolutin. (Up) Cells are in the microfluidic device for 4h before being exposed to an 8min hyperosmotic stress. (Down) Cells are treated with the transcriptional inhibitor for 1h in the microfluidic device, thiolutin is washed out and cells are allowed to grow for 4h before being exposed to an 8min hyperosmotic stress. (D) pSTL1 fluorescence response of cells treated (N=101) or not (N=97) with thiolutin after stress. A similar response is observed. Fluorescence levels are normalized by the value of the peak of fluorescence in the non treated case. (E) Principle of memory effect quantification in the presence of thioluthin. (Up) Cells are exposed to two 8min hyperosmotic stresses separated by four hours. (Down) As as in A, cells are treated with thiolutin for 1h and exposed to a 8min hyperosmotic stress, thioluthin washed out and after a 4h recovery, cells are exposed to second 8min stress. (F) Population quantification of stress memory of cells submitted to a first stress in the absence (blue) or the presence of thioluthin (N=101, pink). Fluorescence levels are normalized by the value of the peak of fluorescence in the non treated case presented in (D). Decrease of fluorescence after second stress is seemingly similar in both cases.

### Chromosome positioning influences the dynamical activity of pSTL1

The STL1 locus is located on the right arm of the chromosome IV, in its subtelomeric region, prone to silencing in non-stress conditions. To investigate the influence of the chromatin context on the dynamics of activation of pSTL1, we moved a region containing the promoter of STL1 and the yECITRINE fluorescent reporter to a distinct, centromeric chromatin domain (figure 4a). The displaced DNA region included 1kb upstream of the STL1 locus, enough to have a fully functional STL1 promoter [33]. To compare pSTL1 activity between its endogenous position and the centromeric one, we first submitted both strains to a 2h hyperosmotic stress and used flow cytometry to quantify the fluorescence at several time points. The activity of centromeric pSTL1 was significantly lower than in wild type cells in two independent clones (figure 4b and supplementary figure 3), although the integrity of the promoter was preserved [33].

**Figure 4.**
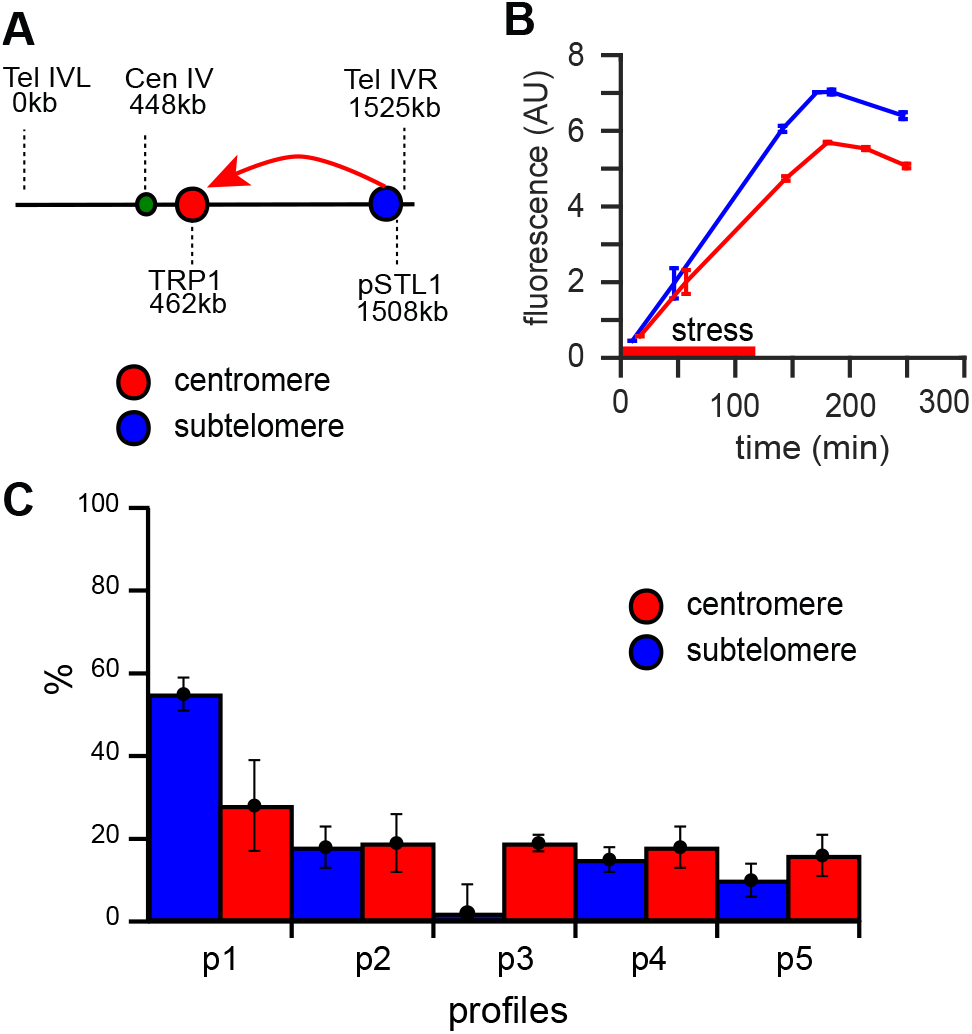
(A) Displacement of pSTL1 towards the peri-centromere of chromosome IV. Sketch of the genomic position of pSTL1 on chromosome IV. The promoter was moved at the TRP1 locus, on the same chromosome. Genomic positions indicated in kb. (B) Decreased activity of the displaced pSTL1 upon stress. Fluorescence quantification of promoter activity in response to a 2h hyperosmotic stress in the endogenous location (blue) and when the promoter was moved (red). Standard deviation of triplicate experiment (C) Displacement of pSTL1 leads to the loss of the memory effect. Single-cell quantification of cells containing displaced promoter pSTL1 in response to two hyperosmotic stresses. Classification made according to the five typical profiles of response as in figure 2.

Patterns of consecutive responses to two 8min hyperosmotic stresses separated by 4h were then compared between endogenous and displaced pSTL1. Cells expressing the STL1 promoter at this centromeric position showed a more uniform distribution into the five defined clusters and there was a decrease in the amount of cells displaying the memory effect (from 55%±11% to 28%±4%, figure 4c) compatible with a purely stochastic process. Such a result indicated that the chromatin environment might be involved in the dynamical transcriptional activity of pSTL1. Although a functional pSTL1 promoter has been displaced, potential subtelomeric regulatory elements could have been lost during gene displacement. We have ruled out this hypothesis using a Crispr/dCas9VPR system [34] to bypass regulatory elements and force the activation of pSTL1 in non-stress conditions (supplementary figure 4). We designed guide RNAs to target Crispr/dCas9VPR within the 1kb sequence of displaced pSTL1 and successfully induced the fluorescent reporter expression independently of any stress at both the peri-centromeric and subtelomeric positions. We observed a decrease in the activity of pSTL1 at the peri-centromeric position compared to the endogenous one, meaning that the observed differences of expression of pSTL1 are not linked to the sequence of the promoter itself or the presence of regulatory elements, but is rather related to the chromatin environment.

Taken together, our results show that the chromatin environment sets the variability of single-cell dynamical response for short stresses and can consequently act on cellular memory.

## Discussion

In the current study, we investigated how individual yeast cells dynamically behave in response to short pulsed hyperosmotic stresses. Focusing our study on short stresses allowed to analyze the genetic response to hyperosmotic stresses exclusively and probe the cell-cell variability that finds its origin in the onset of transcriptional events.

The in-depth single-cell analysis reveals that yeasts display various behaviors in response to two repeated stresses that we have clustered according to several typical profiles. Response to the second stress could be similar, higher or lower than the response to the first stress. The latter case corresponds to the most frequent response and was named memory effect. We have also considered two additional profiles where cells do not respond to one of the two stresses exclusively. These last two profiles might depend on transcriptional delay as verified by simulation and validated by the experimental observation that all cell respond to a long stress, as observed previously [35]. Using stochastic simulations, we have established that the five profiles of response could be explained by gene expression stochasticity. Yet, the single-cell quantification obtained with such a model does not account for the prevalence of the memory effect, indicating it is not a reflection of gene expression stochasticity alone.

Studies on cellular memories in budding yeast in response to repeated stresses typically describe a faster dynamics of gene expression, or lower amplitude of response as two possible ways to respond [5, 7, 36]. Similarly to the response to a long hyperosmotic stress, we observe a decrease in amplitude of pSTL1 response after pulsed stresses, without any difference in the time of reactivation [5]. We speculate that a diminished response could be a strategy for the cells to lessen their burden compared to the case where transcriptional response to two stresses would be similar.

What is the mechanism that could drive a diminished response to a hyperosmotic stress? It is possible that, as observed in the response to hyperosmotic stress triggered by Nacl, a long-lived protein triggered by the stress, could remain in the nucleus upon the first stress and hamper the promoter of interest for a similar activation upon the second stress [5]. However, it is known that after stress, phosphorylated Hog1 translocates within 3 min. into the nucleus, leading to the up regulation of stress response genes, including STL1 and Hog1 rapidly exits the nucleus, 15-30 min after the start of the stress, making this assumption less likely [25]. Alternatively, transcriptional inhibition experiments show that the memory effect does not seem to require de novo protein synthesis. Since transcriptional inhibitor thiolutin inhibits all three RNA Pol and de novo transcription, but does not prevent potential transcription factors from binding to the promoter’s sequence, specific histone marks are possibly left after first stress. Those marks could serve as traces of previous activities of the promoter and could explain a diminished response. Such an interpretation awaits single cell ChIP to be validated.

We further showed that the dynamic variability distribution between single cells was dependent on the positioning of the pSTL1 locus on the chromosome. When displaced in a pericentromeric domain, pSTL1 shows a decreased activity. We could discard potential loss of regulatory sequences by showing that activation of pSTL1 by a CrispR-dCAS9-VPR construct, which bypasses the need for stress factors for the response, still depends on the genomic position. At this pericentromeric position, the stochasticity of gene expression prevails and the memory effect is lost. Consequently, an active mechanism might occur in the subtelomeric regions upon stress, allowing for the memory effect to become predominant among cell population. Interestingly, it has been already observed that changing the position of a gene in the genome can alter its expression [37]. It was proposed that change in expression level could be due to changes in transcription noise, or a in the noisy steps when cells transition between two expression levels [37]. The latter case has been shown to enhance the cellular memory when cells were subjected to glucose limitation stress [20].

One key difference between the two genomic positions we have analyzed is the variation in the amplitude of pSTL1 response. We thus propose that transcriptional marks or transactivators will induce a high level of transcriptional activity during the first stress, required to overcome the stochasticity of gene expression and lead to the emergence of a cellular memory. A parallel can be drawn with previous studies performed in *B. subtilis*, where transcriptional events occurring above a certain threshold have been described to lead to the emergence of a cellular memory [38]. Although this organism is a prokaryote, some similarities might exist between those two microorganisms in regard to the biology of a memory.

It could be hypothesized that a high activity of the promoter of interest could mean an opened chromatin. Although marks of acetylation are usually associated with a high level of gene expression, the activity of pSTL1 is reduced in absence of the histone deacetylase Rpd3 [39]. In the experimental context studied here, marks of deacetylation are possibly involved for a high level of transcriptional activity. It would be interesting to investigate the potential role of (de)acetylation by, for instance, forcing a high level of (de)acetylation during the first stress only.

Stochastic gene expression gives a diversity of behaviors. From an evolutionary point of view, this diversity of responses to repeated stresses allows the selection of the most adapted one. In our experimental conditions, the preference for a memory effect suggests that the specific subtelomeric position of pSTL1 offers an optimal regulation level to perform better adaptation.

Altogether our work shows how critical single-cell studies are for stress memory analyses. It also indicates that establishment and transmission of memory does not require a long stress and can start after short-pulsed stresses. Our work could serve as a basis to broader studies of the positioning of stress response genes in the budding yeast in response to fluctuating environments.

## Materials and Methods

### Flow Cytometry

all flow cytometry experiments were performed with a flux cytometer Gallios (Beckman Coulter) equipped with 10 colors, 4 lasers (488nm Blue, 561nm Yellow, 638nm Red, 405 nm Violet). We used the excitation laser 488nm and the emission filter at 530nm +/− 30 nm.

### Yeast strains and cell culture

our experiments were made using a pSTL1::yECITRINE-His5 (yPH53) strain derived from S288C and gifted to us by Megan McClean. The yeast were grown overnight in SC+2% glucose at 30°C. Cells were diluted next morning to reach OD600=0.5 at the moment of the experiments. Genotypes of all strains used in this study are indicated in table 2. To move pSTL1 reporter construct in the peri-centromeric region of chromosome IV, pSTL1-yECITRINE-HIS5 construct was PCR amplified using primers TATTGAGCACGTGAGTATACGTGATTAAGCACACAAAGGCAGCTTGGAGTCAATGATTCTGAAATACTCCTTTTACA and TGCAGGCAAGTGCACAAACAATACTTAAATAAATACTACTCAGTAATAACATTATTGGTGCGGCAAGG with 50 bases homologies to the TRP1 locus. [HIS+ TRP-] yeast transformants were verified by PCR and PCR fragment subsequently sequenced, ensuring the absence of any mutations in the construct.

### Crispr dCAS9 experiments

we used the plasmid pAG414GPD-dCas9-VPR from Addgene plasmid (# 63801) to express the inactivated form of CAS9 fused to transcriptional activator VPR. The guides were cloned under SNR52 promoter in plasmid pEF534 using the enzyme BsmBI. Digesting the resulting plasmid by NotI / XbaI and cloning the Guide containing fragment into pRS425 similarly digested, performed marker exchange. We designed two guides targeting pSTL1, respectively gRNA1, GAAAGTGCAGATCCCGGTAA and gRNA2, GCGCCGAATACCCCGCGAAA.

### Single-cell clustering

To categorize cells in different classes, we compared the maximum level of fluorescence reached during the first and second stress, while taking the basal fluorescence level prior to the corresponding stress as a reference. The ratio between the maximum amplitude of the first and the second stress was then evaluated. Cells categorized according to profile 1 had a ratio superior to 1, cells categorized according to profile 2 had a ratio inferior to 1 and cells displaying profile 3 had a ratio equal to 1. The cells with a maximum of amplitude equal to 0 (no expression) during the second or the first stress were categorized separately (figure 2b). All ratios were established with a 5% threshold.

### Microfluidics

we used an H-shaped microfluidic device to confine the yeast in channels of 3.7µm high. This microfluidic device was made using soft lithography techniques. The hyperosmotic stresses were triggered using SC+2% glucose supplemented with 1M sorbitol. The media were flown in the microfluidic chip using a peristaltic pump ISMATEC set at 120µL/min flow rate. Mask design is in sup figure 1.

### Transcriptional inhibition

cells were exposed to SC with addition of thiolutin (Abcam ref ab143556) at 50µg/mL (diluted in DMSO) for 1h prior and during the stress. To wash out Thiolutin in the microfluidics experiments, SC medium was delivered to the cells during 4h using a peristaltic pump set at a flow rate of 120µL/min.

### Microscopy

we used an inverted microscope Olympus IX71. Yeasts were observed with an objective ×100 UplanFLN 100×/ 1.3 Oil Ph3 Ul2. Images were recorded with a camera Cool Snap HQ2 Princeton Instruments. All experiments were made at 30°C. Yeast were imaged every 5min with 20ms exposure in bright light and 200ms in fluorescence light. The microscope was controlled by the open source software MicroManager interfaced with Matlab.

## Acknowledgments

Zoran Marinkovic is thanked for help for FACS experiments, Antoine Canat and Pierre Thérizols for helpful advises in thiolutin experiment. The authors would like to thank their respective team members for very fruitful discussions. This work was supported by Labex “Who am I?” (ANR-11-LABX-0071, Idex ANR-11-IDEX-0005-02). EF further acknowledges support from Agence Nationale de la Recherche (ANR-13-BSV8-0013-01), IDEX SLI (DXCAIUHSLI-EF14) and Cancéropôle Ile de France (ORFOCRISE PME-2015). Y.K and E.F. grant support from Fondation pour la Recherche Médicale (ING20160435205). PH acknowledges support by the Agence Nationale de la Recherche (ICEBERG-ANR-10-BINF-06-01) and the European Research Council (ERC) under the European Union’s Horizon 2020 research and innovation program (grant agreement No 724813).

## Author Contributions

ZBM, PH and EF designed the experiments and wrote the article. ZBM performed the experiments. YK, performed Matlab analyses.

## Conflict of interest

Authors declare have no conflict of interest.

## Data availability

All data are available upon request.

---

**Supplemental Figure 1.**
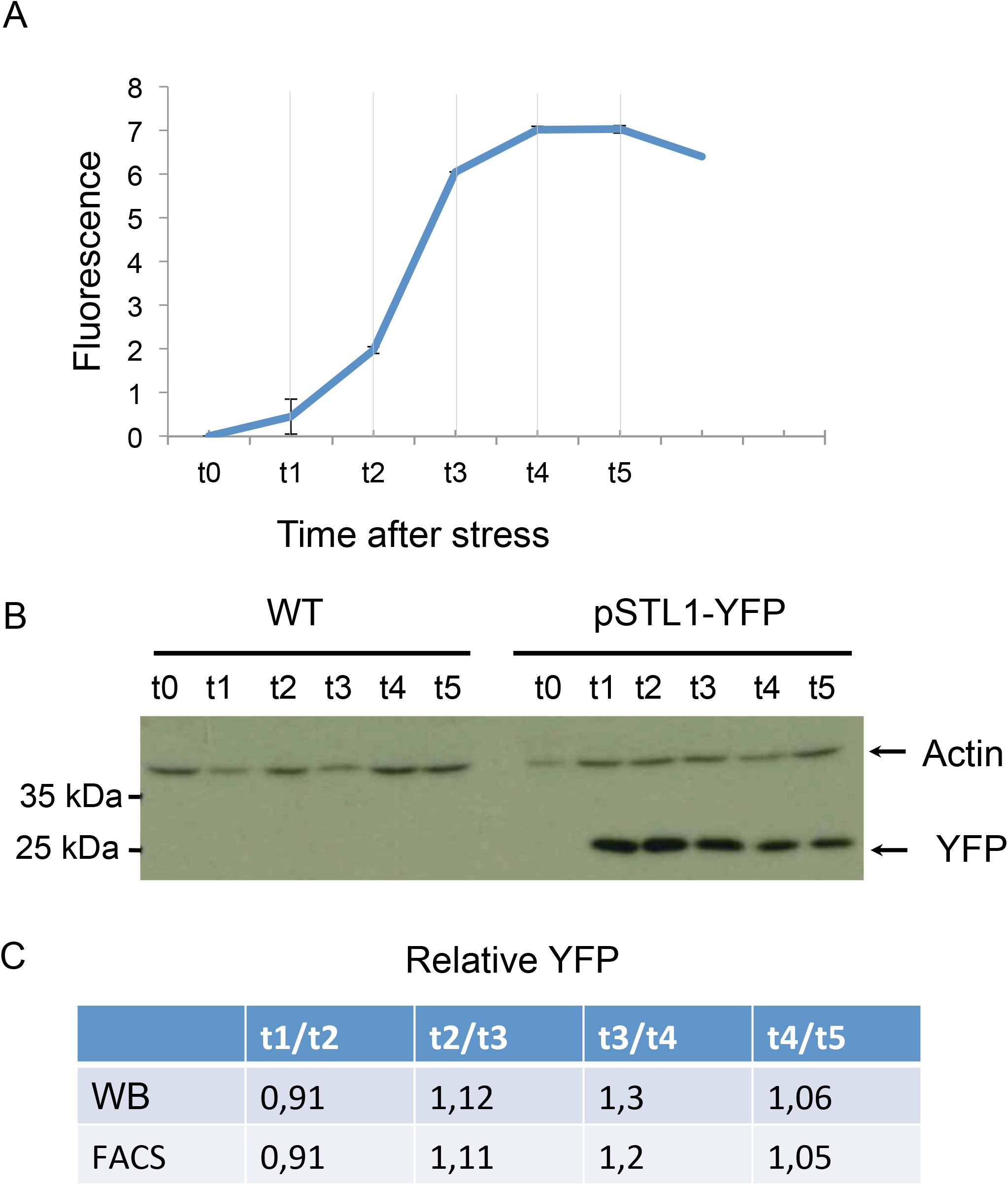
Comparison of relative amounts of pSTL1-YFP by FACs analyses and Western Blotting. (A) After 1h stress in 1M sorbitol, cells were sorted by FACS at different times after stress; 0min (t0), 50 min (t1), 80min (t2), 110 min (t3), 140min (t) and 170 min (t5). Total fluorescence was normalized by fluorescence levels before stress. Note that the minimum of response is reached after 240min. (B) Total protein extracts were performed similarly. Actin was used to normalize the amount of YFP protein expressed. (C). ratios between YFP protein amounts measured by western blot and fluorescence measured by FACS.

**Supplemental Figure 2.**
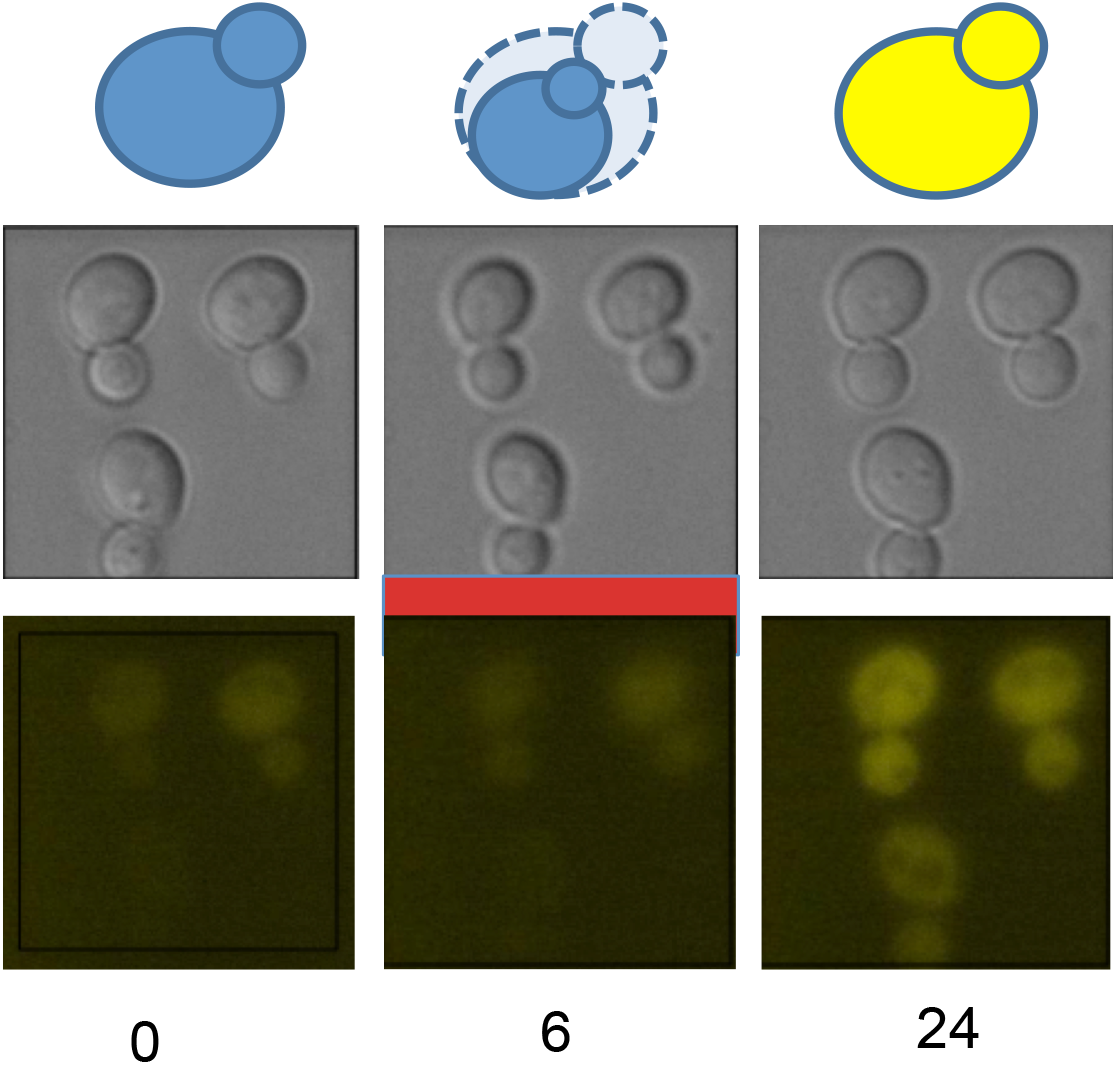
Stress induces cell compression. (A) Example of cells before, during and after stress viewed under nomarski and fluorescence illumination. A scheme of the cells is shown on the top, stress is indicated as a red bar. (B) full video.

**Supplementary Table 1:** Gillespie parameters

| Parameter | Definition | Unit | Reference Value | Source |
| --- | --- | --- | --- | --- |
| k1 | transcription rate | min<sup>-1</sup> | 1.101 | [31] |
| d1 | mRNA decay | min<sup>-1</sup> | 2.94.10<sup>-1</sup> | [31] |
| $\tau$ | time delay | min | Between 0 and 10 min | This study |
| k2 | translation rate | min<sup>-1</sup> | 9.47.10<sup>-1</sup> | [31] |
| d2 | protein decay | min<sup>-1</sup> | 4.10<sup>-3</sup> | [31] |

**Supplementary Table 2:** strains genotype

|  |  |
| --- | --- |
| yPH53<br>(YEF1095) | ura3Δ0 leu2Δ0 his3Δ1 lys2Δ0 pSTL1::yECITRINE-HIS5MX |
| yPH142<br>(YEF1096) | ura3Δ0 leu2Δ0 his3Δ1 lys2Δ0 Δ(pSTL1-STL1)::CaURA3 |
| yPH212<br>(YEF1098) | ura3Δ0 leu2Δ0 his3Δ1 lys2Δ0 Δ(pSTL1-STL1)::CaURA3 Δtrp1::pSTL1-yECITRINE- HIS5MX |
| yPH359 | leu2Δ0 his3Δ1 lys2Δ0 pSTL1::yECITRINE-HIS5 Δtrp1::pURA3-URA3 |

**Supplementary Table 3:**
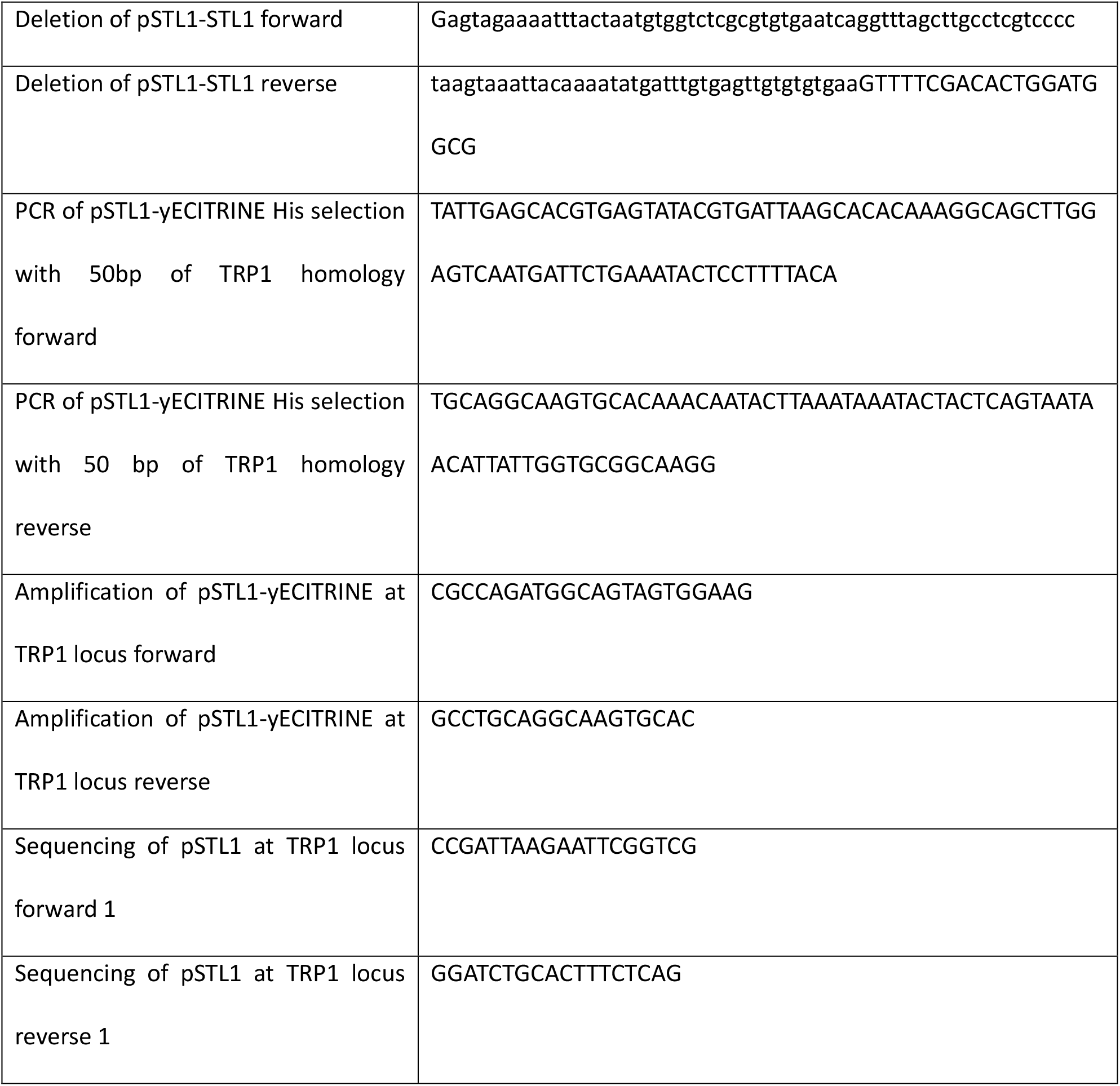
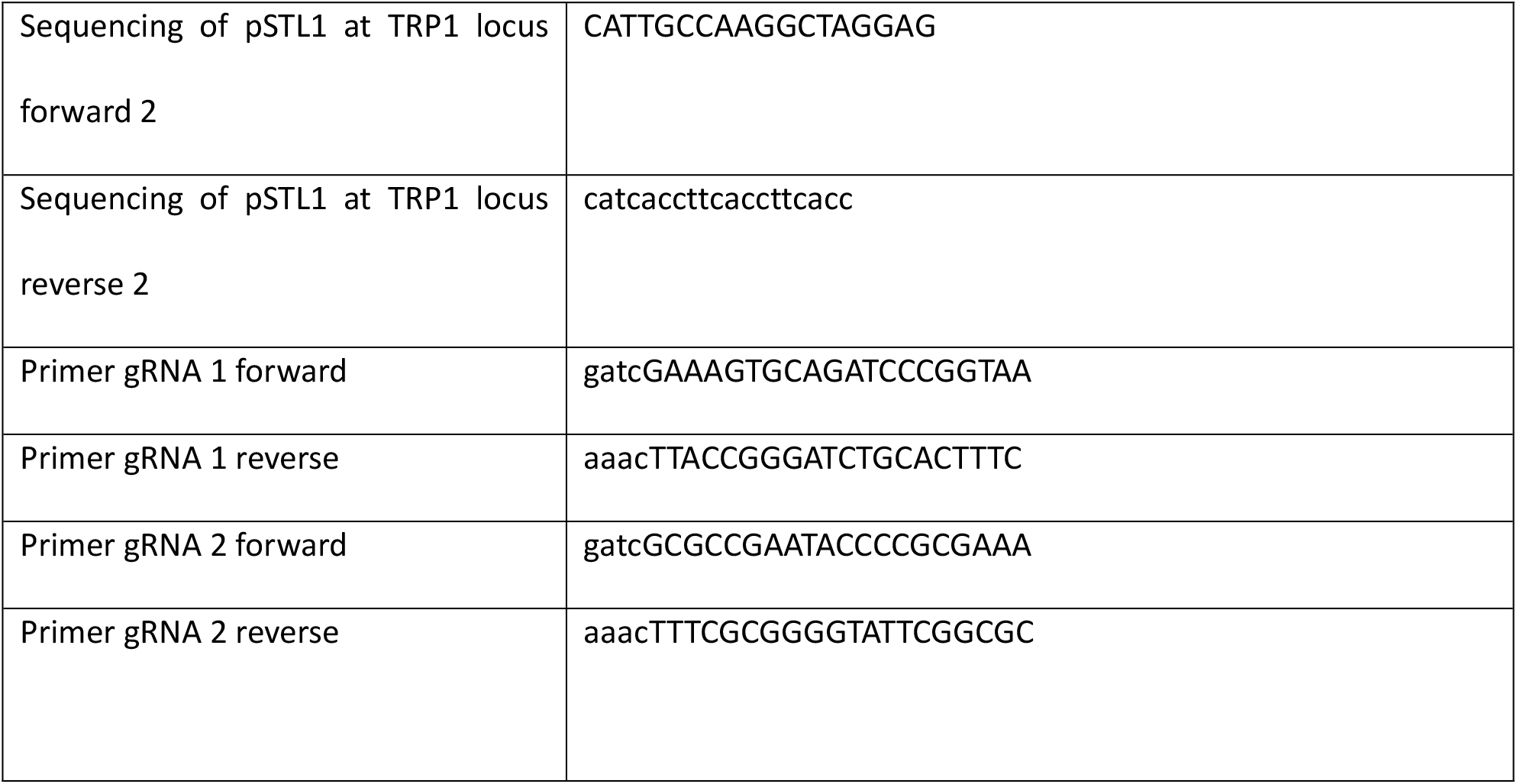
list of primers

